# How goals shape naturalistic memories: bias in gaze and narrative recall

**DOI:** 10.64898/2026.10.28.762515

**Authors:** Hannah Habenicht, Mara Müller, Monika Schönauer, Nora M. Roüast

**Affiliations:** Neuropsychology, University of Freiburg, Freiburg, Germany; Department of Biometry and Environmental System Analysis, University of Freiburg, Freiburg, Germany; BrainLinks-BrainTools, University of Freiburg, Freiburg, Germany

**Keywords:** memory, goal bias, naturalistic design, eye-tracking, semantic segmentation, free recall

## Abstract

Episodic memories are thought to represent information framed by our individual perception and goals, yet personal experiences are hard to evaluate externally. Due to novel methodological advances, we could now trace the development of such goal-shaped memories in a naturalistic experimental setting. We recorded eye-tracking while participants encoded episodic-like memories, and later recorded participants freely narrating their experiences. Naturalistic goal-states were induced by introducing a secret job, which promised to reward participants later for person-based or space-based information. Using computer-vision based semantic segmentation of visual scanning behaviour in movie scenes and data-driven coding of free recall performance, we show that these goals increasingly shift attention during memory encoding and systematically bias narration during retrieval to goal-relevant dimensions. Our findings high-light that we can use data-driven methodology to observe how naturally occurring or induced biases shape how we perceive and remember the world.

## 1. Introduction

Reminiscing about a family holiday, two family members often notice that they came away with different memories of the same situation, perhaps one memory focusing on the tasty food and the other on tense social dynamics. Indeed, even a single person may perceive and remember the same situation differently, possibly motivated at the time by a search for food or social harmony. Such goal states highlight relevant information within our internal and external worlds and this preferential selection by attention leads to prioritised processing (Cowan et al., 2024). The relationship between attention and memory is reciprocal, with selective attention acting on various cognitive levels, such as perception, memory consolidation, and retrieval, yet in turn being guided by existing memories: Prior research has established that both long-term memory or the goal-state in working memory can influence and steer our attention to perceive, interact with, and remember information accordingly (Madore and Wagner, 2022; van Ede and Nobre, 2023). Information deemed goal-relevant, for example due to an association with reward or heightened emotion, is remembered preferentially (Knowlton and Castel, 2022; Dunsmoor et al., 2015). Reward-association also strengthens mnemonic processing of goal-relevant aspects if introduced only after encoding (Patil et al., 2017; Braun et al., 2018), and influences the subsequent consolidation processes during sleep (Sterpenich et al., 2021). The ongoing attentional engagement of the encoding and retrieval of long-term memories can be traced in neural signatures (Song et al., 2021; Sabo et al., 2024) and attention might promote further encoding processes by stabilising mnemonic representations (Aly and Turk-Browne, 2016). Given the evidence that goal-dependent attention biases inform resulting memories and observations of biased memories in daily life, it is highly relevant to characterise how different goal-states shape naturalistic memories and to develop research tools that allow us to trace this process.

If goal-directed attention determines what is remembered, then measuring what is attended to during encoding is a way to anticipate what will endure. Tracking eye movements during naturalistic visual exploration gained traction as a valuable method to assess the deployment of spatial attention, especially given the close links between gaze control, attention, and memory (Jonikaitis and Moore, 2019; Ryan and Shen, 2020). Eye movements during encoding of images predicted resulting memories, whilst gaze during recollection helped judge memory veracity (Damiano and Walther, 2019). Proximity of gaze towards events can index memory, and anticipatory gaze indicates recollection of the associated memory (Schmidig et al., 2025). Additionally, the duration of fixation displayed during scene viewing appears to be the result of memory encoding, not processing demand (Sulewski et al., 2026). Since gaze orients attention selectively, and gaze position can be recorded with millisecond accuracy and high spatial precision, we can track where participants look, for how long, and how often they shift their gaze direction between regions of interest (Rayner, 1998; Kowler, 2011; Duchowski and Duchowski, 2017). Eye-tracking serves as a high-resolution measure of gaze direction, which in turn is directly linked with attention and memory. It may thus provide a promising methodology for investigating how goal-state shapes emerging naturalistic memories.

Whilst eye gaze can serve as a readout of what encoding selects for memory, free recall can provide a naturalistic retrieval measure of episodic memories. Categorised as declarative, episodic memories are typically expressed via language, and free recall therefore resembles natural, self-directed retelling where speakers select which aspects of a remembered event to verbalise (Levelt, 1989). Eye-tracking and free recall measures have previously successfully been paired to trace the encoding and retrieval of the resulting memory, finding that fixation frequency during encoding predicted subsequent free recall across 967 participants (Fehlmann et al., 2020). Celli et al. (2025) reported that the spatial distribution of fixations during free viewing of static images predicted where recalled objects fell in participants’ subsequent verbal reports, and that the strength of this gaze-to-recall correspondence predicted how much each person recalled. What surfaces in spontaneous recall thus carries a trace of what attention prioritised during encoding. Tracking gaze at encoding and free recall the following morning allows us to trace the shaping of memories from first perception to verbal retelling.

Much of this evidence, however, comes from paradigms in which a clearly defined unit, a single scene, an incidental object, or a cued pair, is the entity whose fate in memory is tracked (Damiano and Walther, 2019; Fehlmann et al., 2020; Celli et al., 2025; Sulewski et al., 2026). With advanced methodologies and mechanistic understanding, research can now pivot to emerging complex episodic memories. Episodic memories derive from continuous streams of multisensory information, and recall afterwards is spontaneous and narrative rather than cued item by item. So far, work with eye-tracking in naturalistic film material was limited by methodological constraints: tracking fixations against a moving scene requires knowing which information was presented at which location and was fixated on at each point in space and time. Annotating each pixel, frame by frame, is prohibitively slow for any clip long enough to support a memory task. The powerful readout that gaze fixations offer has therefore often been confined to individual rather than moving scenes. Paradigms that overcome this bottleneck are only now beginning to appear (Trajkovska et al., 2024), and we demonstrate one such approach here using computer vision. Open-source semantic-segmentation models have been developed for tasks such as robotic and automotive vision, to recognise what a camera is viewing from moment to moment. Resulting algorithms can assign a semantic category to every pixel of every frame of a film clip, thus annotating a moving scene automatically. Eye-tracking and automatic segmentation together give a frame-by-frame record of what content a viewer was looking at as the scene unfolded, by mapping gaze continuously onto the semantic content and allowing us to trace attention and encoding in a more naturalistic setting.

Building on these methodological advances, we attempted to trace how emerging episodic-like memories are shaped by goal-related biases using eye-tracking and free recall in a naturalistic task design. We measured unrestricted eye movements while participants watched film clips and later recorded free recall of participants narrating their experiences. To introduce a goal state in an ecologically valid approach, participants were given a cover identity, a Secret Job, that made one category of detail relevant to a rewarded additional mission: personal information for a spy, spatial information for a member of a criminal team. A neutral condition received no such instructions. We predicted goal-relevant information to be preferentially attended to during encoding and recounted during free recall. Whilst humans generally display an attentional bias towards people or faces (Bindemann et al., 2005; Cerf et al., 2009; End and Gamer, 2019; Skripkauskaite et al., 2023), we expected the goal state to modulate the strength of this bias to overemphasise people in the spy condition and depreciate them in favour of spatial information for the criminal condition.

## 2. Methodology

### 2.1. Design

This study aimed to trace how naturalistic memories develop as biased reconstructions shaped by individual goals. The assumption was that goal-aligned information highlighted as important by reward association would receive preferential mnemonic strengthening. These goal states were experimentally introduced with a cover story, a Secret Job, that assigned them a fictional mission focusing on one form of information over others. Spies were told to attend to personal details, people, and faces. Criminals were told to attend to spatial and layout information as well as the structure of the environment. A third, neutral group received no such instruction, and so no goal at all.

Additionally, we wanted to explore how adaptive any shaping of memory formation was, by altering whether the goal state was induced prior to encoding the memories (proactive) or after encoding but prior to sleep-based consolidation (retroactive, Figure 1). A memory system optimised for adaptation might preferentially strengthen goal-relevant dimensions during consolidation even if the goals were introduced only after encoding. Conversely, an adaptive memory system rather oriented towards stability, might focus on strengthening information that was already encoded as goal-relevant. We therefore included five conditions, namely proactive spies, proactive criminals, retroactive spies, retroactive criminals, and a neutral control group.

**Figure 1:**
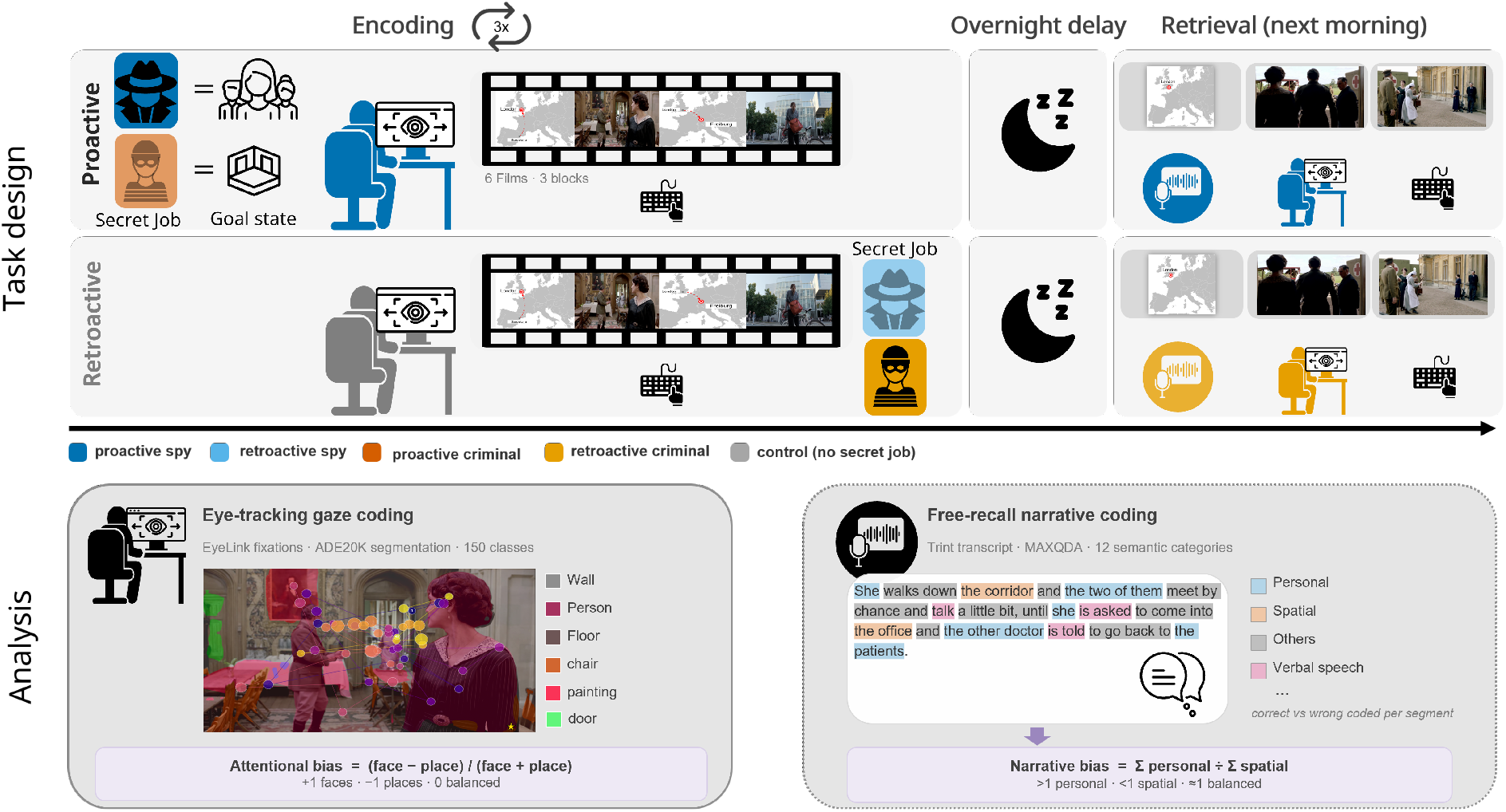
Schematic representation of task design and analysis. **Task design:** Participants viewed six films in randomised order, repeated over three encoding blocks. Experimental groups were assigned a goal state focusing on people (spy, blue hues) or places (criminal, orange hues) either prior to encoding (proactive, darker colours) or subsequently (retroactive, lighter colours). Eye-tracking and behaviour were recorded during encoding. After an overnight delay, participants returned to perform a free narrative recall and a behavioural task during which eye-tracking was recorded. Examples are illustrated for a proactive spy and a retroactive criminal. **Analysis:** Eye-tracking data were coded by pairing gaze location with semantic labels annotated by a computer vision semantic segmentation algorithm. Narrative free recall data were coded by assigning the transcribed words to semantic categories.

The study ran across three phases: an evening encoding session during which goal-states were introduced proactively or retroactively, an overnight delay, and a morning retrieval session. Eye-tracking was recorded during the encoding of naturalistic stimuli in the form of film clips. Retrieval included free narrative recall and behavioural tasks implementing eye-tracking.

### 2.2. Participants

Overall, we examined 100 participants (male: 19, female: 81), aged 18 to 31 (*M* = 22.98,*S D* = 2.89). One participant was excluded from the analyses, given that they fell asleep during the evening session. The remaining 99 participants (male: 19, female: 80, age: *M* = 22.97,*S D* = 2.90) were included in the statistical analysis. After exclusion and due to an assignment error, we were left with a roughly even split across groups (proactive spy: *n* = 19, proactive criminal: *n* = 20, retroactive spy: *n* = 20, retroactive criminal: *n* = 21, neutral: *n* = 19). Inclusion criteria restricted recruiting to right-handed German speakers in the age range from 18 to 35 with normal sight and hearing. Colour blindness, prosopagnosia, as well as sleep disorders led to exclusion. Participants were recruited via the online platform SONA or flyers and were randomly allocated to one of the four experimental groups or the additional control group. Participants were compensated at an hourly rate of 10 Euros. The study was approved by the ethics committee of the University of Freiburg (reference: 21-1603). Informed consent was obtained from all participants. Data collection for this project took place from March through November 2023.

### 2.3. Tasks and measures

#### Naturalistic encoding task, eye-tracking, and engagement task

During encoding, participants watched six short film clips, each roughly 1.5 to 2.5 minutes long, framed as a continuous journey where each film was mapped to a corresponding European city. The film clips were cut from the following German or German-dubbed series or films: Charité (S03 E01), The Intouchables, High Seas (S01 E01), Biohackers (S01 E02), Downton Abbey (S02 E03), and New Amsterdam (S01 E03). The clips provided a naturalistic, dynamic encoding material and we recorded unrestricted eye movements throughout the encoding using an EyeLink 1000 Plus at 500 Hz (monocularly; see Section 2.6.1 for the full processing pipeline).

While watching, participants performed a continuous target-detection task: coin icons (50×50 pixels) appeared at predefined locations within each clip, and participants pressed a key as soon as they spotted one. We placed these coins within content from one of three semantic categories: personal, spatial, or object. This engagement task encouraged participants to view the entire film display irrespective of goal, in order to make experimental conditions more comparable. Furthermore, the detection rate of coins also served as a behavioural index of general learning of the encoding material, and of category-specific allocated attention through reaction time and hit rate. Increasing hit rates would indicate greater familiarity with the film material.

#### Retrieval tasks

The next morning, we assessed memory for each of the six film clips first via free recall and then via a coin replacement task, in randomised order. Only once tests for all six cities were completed, did participants receive a reminder of their goal-state and completed the secret job task, namely a rewarded goal-relevant multiple-choice test. Since it contained only a single trial, the latter was not statistically analysed.

#### Free recall

The map interface from the journey through Europe introduced each city, and participants were asked to report everything they could remember about the events shown in that clip verbally and in as much detail as possible. No reference to their secret job was made, nor were specific retrieval cues given. Participants could terminate their recording themselves, or it timed out after the maximal time of four minutes per city. Free recall complemented eye-tracking during encoding as a naturalistic retrieval method. Given the absence of cues, any goal-consistent shift in what participants chose to report reflects what survived in memory and was selected for recall.

#### Coin replacement task

Participants were presented a screenshot from the original clip and asked to replace a coin where it had appeared during encoding. In addition to reaction time, memory precision was quantified as the Euclidean distance between the placed coin and its true original location. Apart from confirming large precision differences across personal, spatial, and object categories, we could not analyse this task further due to low task adherence.

### 2.4. Procedure

After providing informed consent, participants were given a vision test to determine eye dominance. They were seated at the computer and received written and oral task instructions. Participants also completed a short computer-based practice task, which additionally tested the headphones and volume. The eye-tracker was calibrated to their dominant eye. Participants then completed all three blocks of the encoding task, with film order randomised for each block. Depending on random assignment, they were either informed of their secret job identity before the encoding task (proactive), after it (retroactive), or received no information regarding secret jobs (control). Secret job information was only given via the pre-recorded audio-visual clip on the screen, and was neither addressed nor discussed by the experimenters, along with the idea of it being secret. Participants then returned home to sleep for the night.

The following morning, participants returned to the laboratory. Participants again received written and oral task instructions as well as a computer-based practice task. As part of the practice, a short narrative was recorded to test the microphone integrated in the headphones. After the eye-tracker calibration, participants completed the free recall and coin replacement tasks for each film clip. The order of films was randomised across participants. After completion, participants alerted the experimenter, who explained and started the final task termed Secret Job task. Finally, after the rewarded secret job task, participants filled out their payment information and were debriefed on the study.

### 2.5. Technical equipment

For encoding and retrieval tasks, participants sat in front of a 24-inch iiyama monitor (ProLiteGB2488HSU) with a spatial resolution of 1920 × 1080 pixels and a refresh rate of 144Hz, and connected to the Host Pc (Dell Optiplex 7010). An EyeLink 1000 Plus eye-tracker was used to track gaze behaviour at 500 Hz using monocular tracking. Calibration and validation were conducted prior to the task (9-point procedure; 5-point if necessary). Head position was fixed using a height-adjustable chin rest, positioned 80 cm from the screen centre and 60 cm from the eye-tracking camera. Behavioural output was measured via a keyboard with a German key layout. Sounds were presented and recorded via a headset with a microphone by Logitech. All tasks were programmed and presented with PsychoPy (version 2022.2.4).

### 2.6. Data processing and analyses

Data processing and analyses were preregistered at AsPredicted.

#### 2.6.1. Eye-tracking data

A raw gaze coordinate on its own tells us only where on the screen someone looked, not what was presented at that location. Since our film clips are dynamic, the content at any given screen position changes from frame to frame, so relating gaze to the categories that mattered – personal, spatial, object – required knowing exactly what occupied a point on the screen in any given frame (Figure 1).

We converted the raw output by the EyeLink 1000 Plus from EDF to ASCII format using SR Research’s Data Viewer and parsed the converted files with custom Python scripts. EyeLink’s built-in velocity and acceleration criteria classified each recorded sample as part of a fixation, a saccade, or a blink; fixations formed our primary unit of gaze behaviour, with the continuous sample stream retained as a validation measure. We split each recording into chunks corresponding to individual film clips using the clips’ onset and offset timestamps, and matched every gaze timestamp within a chunk to its nearest video frame based on frame rate (Supplementary Figure A.5).

To identify what a participant was looking at, we passed each video frame through a pre-trained convolutional neural network (CNN), the ADE20K-ResNet101-UPerNet semantic segmentation model (Zhou et al., 2017, 2018), which labels every pixel in a frame with one of 150 categories. For each fixation, we read off the label at the matching pixel in its onset frame; the continuous sample stream was labelled the same way (Supplementary Figure A.5). We collapsed the 150 ADE20K categories into our four theoretically defined classes, personal, spatial, object, and other, matching the Secret Job categories (see Appendix A.2) and the coding scheme used for the memory analyses. This gave us a semantic category for every gaze sample during encoding and retrieval. The primary outcome of the eye-tracking analysis was therefore the distribution of these categories across the viewing period: our index of goal-consistent attentional bias, measured at the moment it could first have arisen.

For each film viewing, we classified fixations (and, in parallel analyses, dwell time and raw gaze samples in 500 Hz) by the semantic region gaze landed on, and excluded observations that fell outside of the monitor boundaries (1080×1920 px). Using these measures, we quantified where participants looked, the distribution of semantic categories viewed, the number of fixations and their duration (ms), and visual exploration behaviour via the scanning path in pixels. Additionally, attentional bias toward faces versus places during encoding was formalised as a normalised bias score, *B* = (face−place)/(face+place). Objects and uncategorised regions were excluded from the denominator. *B* ranges from −1 (exclusive attention to places) through 0 (equal attention) to +1 (exclusive attention to faces). We computed the score separately for three convergent gaze measures: fixation count, fixation duration, and raw gaze-sample count.

Beyond missing files, a subset of recordings showed recording and calibration issues: fixations which clustered implausibly at the top of the screen, pointing to calibration drift rather than behaviour. We added a participant-level tracking-quality gate, combining experimenter comments logged during the session with a quantitative check on the raw gaze stream: for each participant, we computed the proportion of fixations and sample timestamps falling outside the screen area (out-of-bounds, OOB) across the whole recording. Participants with an OOB rate at or above 10% on either measure were excluded, as were those already removed for missing files or task non-compliance. This gate flagged 11 participants, 7 of whom were not already excluded on other grounds, so that 16 participants were excluded in total, leaving 79 participants in the encoding eye-tracking sample.

##### Statistics

Each film viewing was assigned to the proactive spy or proactive criminal group (role instruction given before the film) or to a Control group collapsing all remaining viewings (retroactive and neutral conditions). Since retroactive instructions could not influence encoding gaze, retroactive conditions were treated as equivalent to control.

For each gaze measure, bias scores were analysed with a linear mixed-effects model:

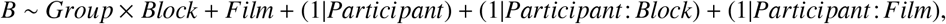

with Group, Block (three viewing repetitions) and Film as fixed effects. Film was included as a fixed effect because the six clips differed in their face/place content. The random intercepts accounted for the repeated-measures design: participants could differ in their overall bias, how their bias changed across blocks, and in their individual bias towards the films. Models were fitted using restricted maximum likelihood, with Satterthwaite-approximated degrees of freedom. Group differences were tested with Tukey-corrected pairwise contrasts of estimated marginal means (Proactive spy vs. Control, Proactive criminal vs. Control, and Proactive spy vs. Proactive criminal), reported with 95% CIs and Cohen’s *d*, standardised by the total SD (all random-effect and residual variances plus the variance of the film effects). Estimated marginal means for each group were additionally tested against zero (Bonferroni-corrected for three tests) to establish whether that group showed a face bias at all, independently of the between-group contrasts above.

#### 2.6.2. Behavioural data

Response data to the coins appearing during the encoding phase were analysed on a trial-by-trial basis with regard to detection of and reaction time to the coin appearance. Data were cleaned based on response time deviations beyond three standard deviations of the subject mean by spatial category and block, and an absolute cut-off for response times beyond 2.5 seconds. Outlier detection across participants for accuracy beyond three standard deviations of the mean confirmed a catastrophic drop-off in performance for later encoding blocks for one participant, who was subsequently excluded from all analyses as eye-tracking measures were consistent with the participant falling asleep. Retroactive conditions were again treated as control groups during the encoding phases.

Detection accuracy, with two possible outcomes of detected (1) or not (0), and log-transformed response time measures for detected trials were both analysed on the individual trial level. For this purpose, we fitted a generalised linear mixed-effect model using a binomial distribution with logit link and a linear mixed-effect model, respectively, using the lme4 package in R. The chosen model used Category (spatial placement of the coin: face, place, object), Group (condition: spy, criminal, control), and Block (three viewing repetitions) as fixed effects and Film as a fixed control factor to remove systematic between-film differences. The random-effects structure comprised random intercepts and random slopes for category by participant, and random intercepts for the participant by film interaction:

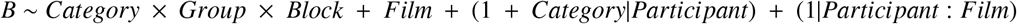

We selected this structure because the maximal specification did not converge, and block slopes instead of category slopes made negligible differences. Type III tests were obtained with Wald *χ* tests (accuracy) and *F* tests with Satterthwaite’s method (RT) as well as Tukey-corrected post-hoc pairwise comparisons for significant main or interaction effects.

#### 2.6.3. Free recall data

For each participant, six audio files were created (.wav) corresponding to the six film clips. The file names were condition-blinded and the audio files were uploaded to the transcription software Trint for automated transcription. Quality control of the resulting transcripts was performed manually, and the combined transcripts were linguistically analysed using the linguistic programme MAXQDA (versions 22.8.0 and 26.1.0). For the linguistic analysis (Figure 1), we predefined twelve semantic categories in accordance with goal-relevant and control categories, adapted from (Rädiker and Kuckartz, 2019): personal details, wrong personal details, spatial details, wrong spatial details, object details, wrong object details, verbal speech details, wrong verbal speech details, not movie related, signs of insecurity, others and wrong others. The detailed category definitions can be found in the supplementary documents (Appendix A.2). Each word in the transcript was assigned to one of these categories by a condition-blinded rater. This allowed separating the narrative content of the free recall in goal-relevant and irrelevant dimensions as a percentage of the number of overall coded segments. The output also allowed controlling for whether the recounted details were factually accurate or not, or combining them into an overall dimension.

Narrative behaviour was quantified by overall word count and the percentage of words dedicated to the semantic categories (namely, spatial, person, object, speech, unrelated, insecurity, or other). In order to assess whether narrative content was biased towards personal or spatial information, we additionally calculated a narrative bias score by dividing the percentage of personal segments by the percentage of spatial segments. A bias score over one indicated a general narrative bias towards personal details, below one towards spatial details, and around one of a balanced representation of personal and spatial details. A normalised bias measure analogous to the eye-tracking analysis (2.6.1) was also calculated as additional exploratory analysis.

##### Statistics

Statistical models were calculated for the same subgroups, first considering only proactive and control, then only retroactive and control groups (*B* ∼ *Group*), before contrasting proactive and retroactive groups for the narrative bias score (*B* ∼ *Timing x Type*). For the word count, we fit a negative binomial model using likelihood-ratio as omnibus tests, for the percentage/proportion values a beta regression with logit link using likelihood-ratio tests, and for the narrative bias scores OLS linear regression models using Robust Wald F tests and planned contrasts with robust standard errors for post-hoc tests to account for non-normal distributions. Given missing free recall data (2) and the excluded participant based on falling asleep during encoding (1), these analyses included a total of 97 participants. For the narrative bias score analysis, a further participant (retroactive spy) had to be excluded due to being a statistical outlier, even though the exclusion did not change the pattern of results.

#### 2.6.4. Attention and narrative bias correlation

Spearman correlations between the attention and narrative bias scores were calculated using either the mean attention bias score across the three encoding blocks, or the modulation of the attention bias by calculating the difference between the first and last encoding block (*B* = Block 3 − Block 1). The attention bias was drawn from each of the three eye-tracking metrics, namely fixation count, fixation duration, and raw sample proportion.

## 3. Results

### 3.1. Naturalistic viewing behaviour modulated by goals during encoding

We first analysed viewing behaviour during the encoding period to establish the general pattern of gaze across participants: where they looked, how this developed across blocks, and how it differed between groups (Figure 2A). Of the 100 participants tested, five had no usable eye-tracking recording and a further 16 were excluded for eye-tracking quality reasons, leaving a reduced sample for the encoding eye-tracking analyses (*n* = 79).

**Figure 2:**
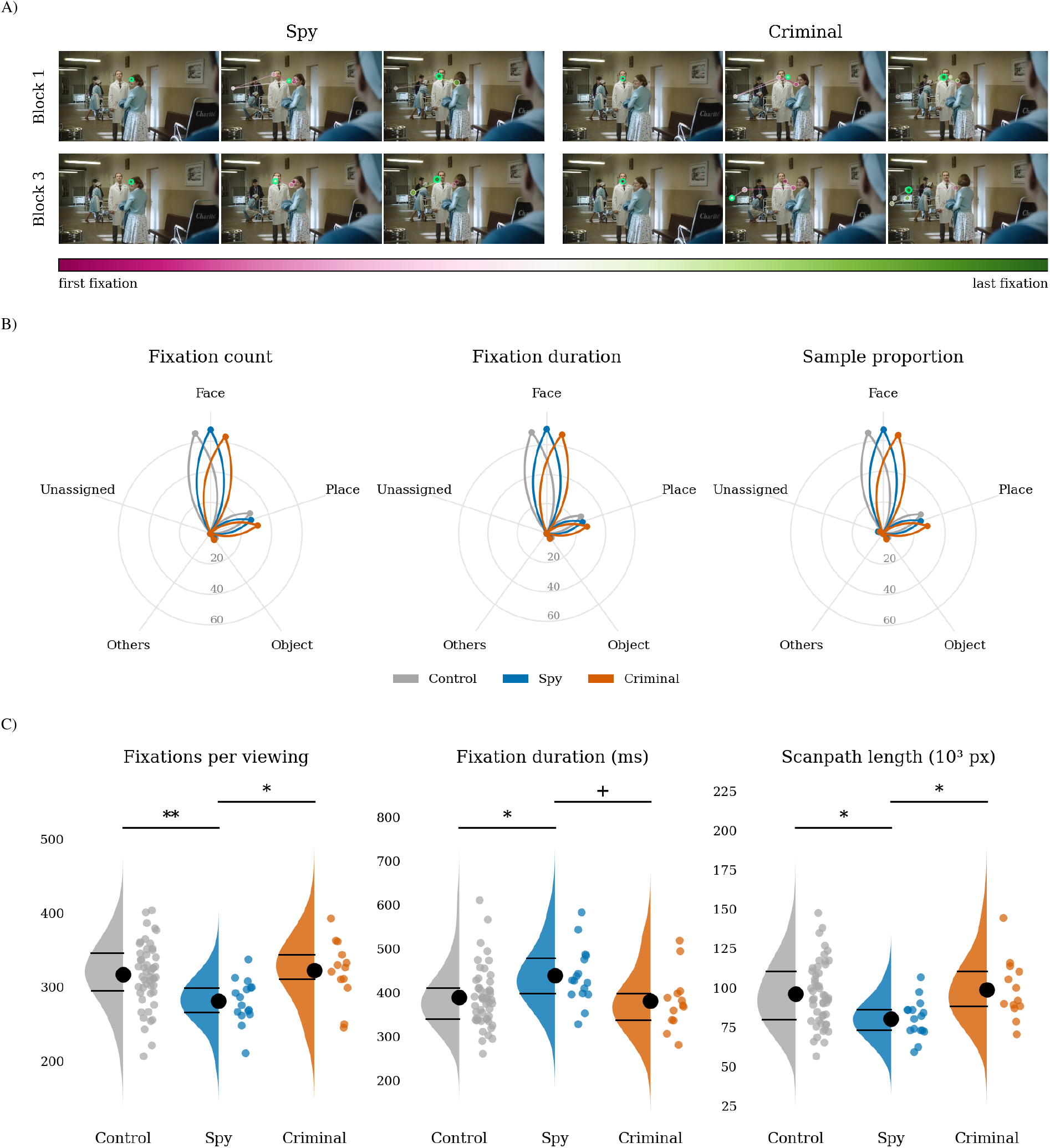
Where participants look. **A.** Time-resolved gaze filmstrips for two participants viewing the same scene from Charité, contrasting viewing behaviour between encoding condition (Proactive Spy vs. Proactive Criminal) and repetition (block 1 vs. block 3). Colour encodes eye-tracking fixation order within the scene with the green circle representing the current fixation during the frame. Circle radius is proportional to fixation duration, min–max scaled to the longest fixation within that scene. **B.** Radial plots illustrate the percentages of semantic content viewed across conditions for fixations, duration, and sample proportions. Across all conditions, most viewing time was allocated to faces, followed by spatial information. Colours denote encoding condition (control: grey, spy: blue, criminal: orange). Solid line indicates mean % with shaded bands = mean ± SEM, averaged across blocks & films. **C.** Violin plots illustrate differential viewing behaviour shown by proactive spies in contrast to criminals and control groups, showing a decreased number of fixations, yet held for longer durations, resulting in an overall decreased scanning path length in pixels.

#### Where people look

From the recorded eye-tracking data, fixation locations were counted for each semantic category across the three encoding blocks. Across every condition, participants allocated most of their viewing time to faces, followed by spatial information, with objects receiving comparatively little attention. The same pattern was apparent for fixation count, fixation duration, or the raw proportion of gaze samples (Figure 2B). On average, faces drew the largest share of valid viewing time (64–71% across the three measures), places the second largest (25–31%), and objects markedly less (around 4%). The residual ‘others’ category accounted for well under 1% of viewing time throughout, whilst raw gaze samples carried an additional 2–4% share left unassigned due to blinks and lost tracking (see 2.6). A within-participant Friedman test comparing the four categories returned *χ*^2^(3) = 237.00, *p* < .001 for fixation count, fixation duration, and sample proportion alike - every one of the 79 participants ranked the categories face *>* place *>* object *>* others identically, whichever measure was used. This ranking held within every instruction group, but its scale did not: proactive criminals gave faces a smaller share of their viewing time than control or proactive spies (fixation-count face share: 63.8% against 66.2% and 67.8%), foreshadowing the face-place dissociation formally tested in the next section (see 3.2).

#### Search breadth

Additionally, we assessed further metrics of gaze behaviour across participants: how many fixations were made per viewing, how long each fixation lasted, and how far their gaze travelled between fixations. These measures were independent of the semantic content fixated on and were used for exploratory analyses based on the observation that proactive criminals seemed to explore the scene more freely than spies. We further computed the spatial dispersion of fixations around their own centroid, a measure of how much of the screen participants covered rather than how they moved across it. Each measure was analysed with a linear mixed model with fixed effects of group, block, and their interaction as well as the same film and random-effects terms as the bias model (see 2.6.1).

We found that group condition significantly affected the number of fixations per viewing (*F*(2, 76) = 5.44, *p* = .0062), mean fixation duration (*F*(2, 76) = 3.86, *p* = .025), and scanpath length (*F*(2, 76) = 4.96, *p* = .0094), but not dispersion (*p* = .310). In every case, the pattern was the same: proactive criminals resembled control groups (all Control-Criminal contrasts: Tukey *p* ≥ .89, |*d*| ≤ 0.13), whilst proactive spies departed from both other groups. Spies made fewer fixations (*M* = 282, *S E* = 10.1) than control (*M* = 318, *S E* = 5.7; *t*(76) = 3.10, *p* = .0075, *d* = 0.66, 95% CI [0.24, 1.09]) or criminals (*M* = 323, *S E* = 11.2; *t*(76) = −2.72, *p* = .022, *d* = −0.76, 95% CI [−1.31, −0.20]).

Fixation durations also lasted longer for spies (*M* = 439 ms, *S E* = 16.8) than for control (*M* = 389 ms, *S E* = 9.5; *t*(76) = −2.60, *p* = .030, *d* = −0.62, 95% CI [−1.10, −0.15]), with a comparable contrast against criminals that reached only trend level (*M* = 382 ms, *S E* = 18.6; *t*(76) = 2.31, *p* = .060, *d* = 0.72). Finally, gaze travelling length was shorter for spies (*M* = 80,592 px, *S E* = 4690) than controls (*M* = 96,437 px, *S E* = 2660; *t*(76) = 2.94, *p* = .012, *d* = 0.73, 95% CI [0.24, 1.23]) or criminals (*M* = 99,064 px, *S E* = 5210; *t*(76) = −2.63, *p* = .027, *d* = −0.85, 95% CI [−1.50, −0.21]). This pattern matches what we would expect from narratively driven film clips, where people tend to be centred within the camera’s focus, meaning that a viewer fixating on faces has less need to look further afield. Spies could satisfy their goal-relevant category, people, with fewer, longer, centrally anchored fixations, whilst criminals, whose category was the surrounding space, had no such shortcut and scanned like control (Figure 2C).

These measures of viewing behaviour were not independent because they used the same film material and thus shared the same duration: longer fixation durations leave less time within that window for further fixations, and fewer fixations mechanically shorten the total distance travelled between them. Fixation count, duration, and scanpath length therefore describe one underlying strategy from three angles: spies dwelt longer on each fixation and, as a direct consequence, made fewer of them and covered less distance overall. The dispersion measure isolated coverage from pacing, since it assessed how widely fixations were scattered across the scene, rather than how many fixations occurred or how far apart they fell in time. The absence of a condition difference for dispersion overall and within every block (*p* ≥ .30, all |*d*| ≤ 0.32, all confidence intervals crossing zero), indicates that spies reached the same visual ground as control and criminals. Goal-state did not change the extent of visual ground covered, but the pace at which they moved across it.

When examining the learning and habituation across blocks, we found that the fixation-count and scanpath effects were already present or trending in the first viewing block and grew more reliable with repetition (scanpath Control-Spy: *p* = .033, *p* = .014, *p* = .008 across blocks 1-3, Tukey-adjusted; fixation count Control-Spy: *p* = .055, *p* = .004, *p* = .006 across blocks 1-3, Tukey-adjusted), yet the group × block interaction itself fell short of significance for either measure (fixation count: *F*(4, 152) = 0.92, *p* = .457; scanpath: *F*(4, 152) = 0.61, *p* = .656). The pattern was similar for fixation duration, with no significant interaction between condition and block (*F*(4, 152) = 0.99, *p* = .416), yet a widening Control-Spy gap across the session, from a non-significant 40 ms in block 1 (*t*(97.8) = −1.95, *p* = .131) to 48 ms in block 2 (*t*(97.8) = −2.33, *p* = .056) to 62 ms by block 3 (*t*(97.8) = −3.03, *p* = .009), whilst control and criminal durations stayed flat. Across repeated exposure to the same film, spies continued to make fewer fixations and their gaze travelled less far, and yet they tended to increase how long they dwell on the fixations.

### 3.2. Goal-state shaped the attentional bias score towards goal-relevant categories

Since the proactive spy and criminal roles were built around goal-relevant categories, spies tracking a rogue contact and criminals surveying a location, we expected spies to focus more on people and criminals on spatial detail across scenes (Figure 3A). To quantify whether goal-state induced such a differential viewing pattern, we analysed the normalised attentional bias score based on fixation data. Positive values indicated a bias towards faces and negative values a spatial preference. The bias score was analysed with a linear mixed model using fixed effects of group, block, and their interaction, controlling for film and with random intercepts for participant, participant-by-block and participant-by-film (see 2.6.1).

**Figure 3:**
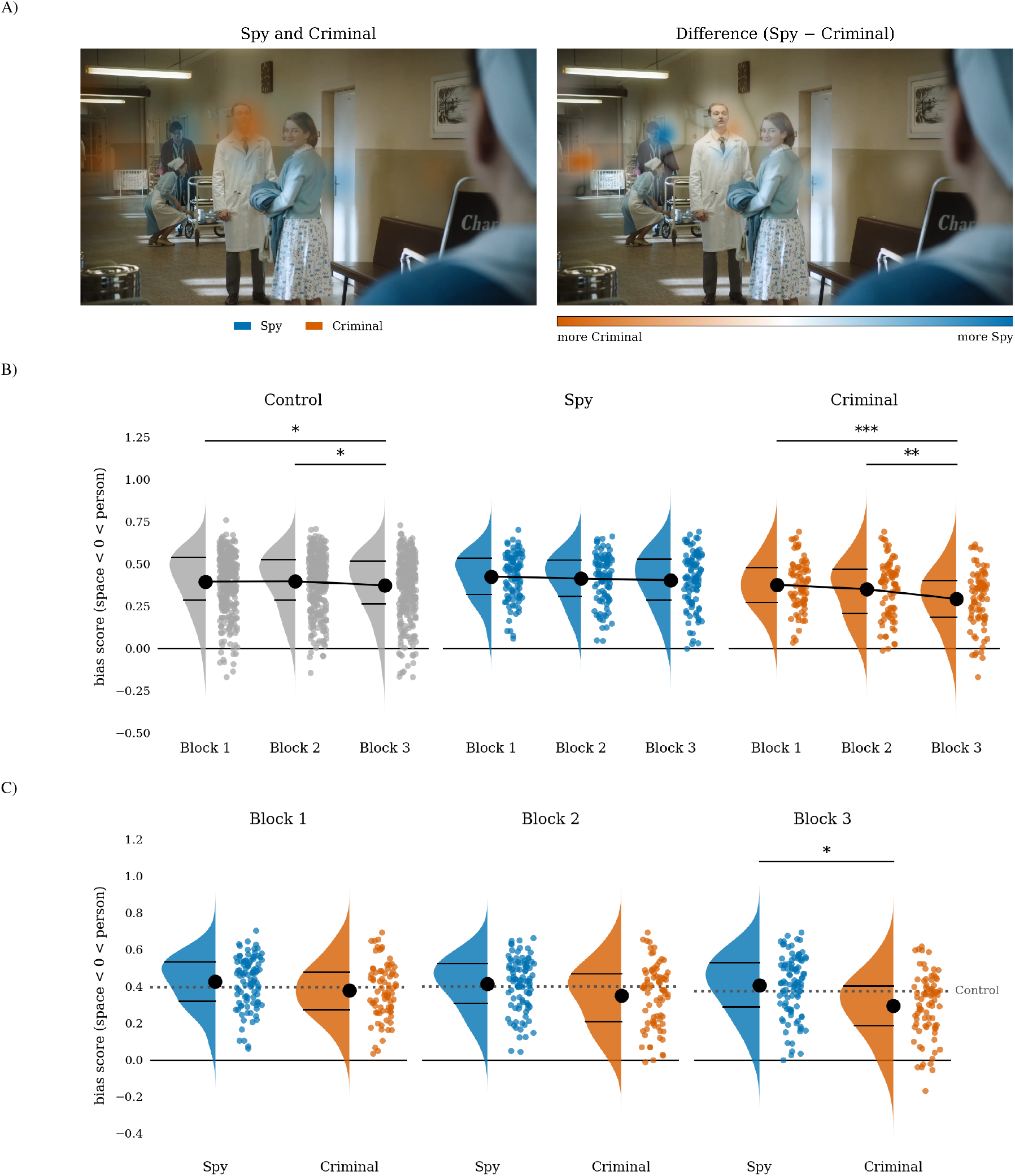
Goal state modulated viewing bias. **A.** Group differences in gaze allocation when viewing a Charité scene in the final encoding block. The left-hand panel shows fixation density for proactive spies (blue, n = 15, 72 fixations) and proactive criminals (orange, n = 13, 62 fixations) overlaid on the scene frame; density is duration-weighted and averaged per participant, overlaid on one brightness scale. The right-hand panel shows the mathematical difference of same two densities (Proactive Spy – Proactive Criminal) on a diverging blue–white–orange scale, isolating where the gaze diverges by condition with colour saturation indicating greater divergence. Blue hue indicates more spy gazes, for example on the male background character, orange hue indicates more criminal gazes, for example on the door frame in the background. **B.** Violin plots show the development of the normalised attention bias across blocks for the different encoding conditions. Bias score here shown for the fixation-count measure, rescaled −1 to +1, with positive numbers indicating greater person bias and negative numbers greater spatial bias. Whilst all groups showed a marked person bias, the bias score decreased slightly across blocks for control groups (grey), stayed markedly high for spies (blue), and decreased most strongly for the criminal group (orange). **C.** Further violin plots illustrate the growing difference in bias score between spy and criminal conditions shown in B., by contrasting them for each encoding block with the mean of the control group as reference line (dotted, grey).

As already seen in raw viewing behaviour, every group showed a significant positive face bias throughout encoding (Figure 3B): proactive spies fixated faces more than places (*M* = .42, *S E* = .03, *p* < .0001, *n* = 16), as did proactive criminals (*M* = .34, *S E* = .03, *p* < .0001, *n* = 13) and controls (*M* = .39, *S E* = .02, *p* < .0001, *n* = 50; Bonferroni-corrected for three tests). The absence of a significant group effect (*F*(2, 76) = 1.61, *p* = .208) indicated that all groups showed a comparably strong bias towards faces. Indeed, a planned post-hoc test confirmed that proactive spies and proactive criminals did not differ significantly from each other when averaged across blocks (*t*(76) = 1.77, *p* = .188, *d* = 0.43, 95% CI [−0.06, 0.92]).

However, viewing behaviour changed across the encoding session, as indicated by a significant effect of block (*F*(2, 152) = 16.51, *p* < .0001). Face bias fell across the session: from block 1 (*M* = .40) to block 3 (*M* = .36; block 1 vs. 3: *t*(152) = 5.60, *p* < .0001; block 2 vs. 3: *t*(152) = 3.92, *p* = .0004). Importantly, there was a significant interaction between group and block (*F*(4, 152) = 3.46, *p* = .0098), indicating that the goal-state influenced how gaze behaviour developed across the session (Figure 3B). As post-hoc tests confirmed, proactive spies maintained their viewing bias towards faces across blocks, as incentivised by their goal state, showing no significant change even across the largest difference between the first and third block (block 1 vs. 3: *t*(152) = 1.53, *p* = .278, Tukey-corrected). In contrast, participants in the control condition showed a continuous decrease in the bias score away from faces (block 1 vs. 3: *t*(152) = 2.75, *p* = .018; block 2 vs. 3: *t*(152) = 2.91, *p* = .011). Crucially, proactive criminals with a goal state motivating a viewing bias away from faces showed not only a decrease in the bias score away from faces and towards spatial information but this shift was more pronounced than in the control condition (block 1 vs. 3: *t*(152) = 5.27, *p* < .0001; block 2 vs. 3: *t*(152) = 3.61, *p* = .0012). These results indicate that the goal-state modulated the naturally occurring decrease in face bias, both by maintaining it in the spy condition and by emphasising it in the criminal condition.

We further calculated between-group contrasts to confirm that the development of the viewing bias differed according to the assigned goal state (Figure 3C). The Control-Spy contrast stayed flat and small throughout (Block 1: *t*(89.2) = −0.90, *p* = .643; Block 2: *t*(89.2) = −0.46, *p* = .888; Block 3: *t*(89.2) = −0.91, *p* = .637; Tukey-adjusted). The Control-Criminal and Spy-Criminal contrasts widened steadily: Control–Criminal grew from Block 1 (*t*(89.2) = 0.53, *p* = .858) through Block 2 (*t*(89.2) = 1.28, *p* = .410) to a trend by Block 3 (*t*(89.2) = 2.20, *p* = .076, *d* = 0.47, 95% CI [0.05, 0.89], Tukey-adjusted; uncorrected: *p* = .030), and Spy–Criminal grew from Block 1 (*t*(89.2) = 1.13, *p* = .498) through Block 2 (*t*(89.2) = 1.43, *p* = .332) to a significant difference by Block 3 (*t*(89.2) = 2.53, *p* = .035, *d* = 0.65, 95% CI [0.14, 1.15], Tukey-adjusted). By the last viewing block, proactive spies showed a significantly stronger face bias than proactive criminals, a gap not yet present earlier in the session. This effect seems to be partially driven by the depreciation of faces in the criminal condition, given the emerging difference of criminals to controls. Furthermore, testing interaction contrasts by evaluating the growing difference between blocks 1 and 3 confirmed that the separation was driven by the criminal condition in contrast to controls (*t*(152) = 3.45, *p* = .002; Holm-adjusted). These results confirmed a modulation of viewing behaviour by goals: Whilst a natural bias towards faces decreased across blocks, this decrease was diminished by the spy goal-state and enhanced by the criminal goal-state.

Both alternative measures reproduced this pattern. We recomputed the bias score from fixation duration (the total time fixations dwelt on each region) and from the raw sample proportion (the share of gaze samples falling on each region, independent of fixation detection). Both confirmed the group × block interaction (raw sample proportion: *F*(4, 152) = 3.00, *p* = .020; fixation duration at trend level: *F*(4, 152) = 2.22, *p* = .069). The Block 3 spy–criminal dissociation held for the raw sample-proportion score (*t* = 2.49, *p* = .038) but was not significant for fixation duration (*t* = 1.90, *p* = .146). However, for both measures, criminals depreciated faces across blocks more strongly than controls (sample proportion: *t*(152) = 3.23, *p* = .005; fixation duration: *t*(152) = 2.66, *p* = .026; Holm-adjusted).

To validate the composition of the control group, we compared the two retroactive groups, who received their role assignment only after encoding and so carried no goal state while viewing, against control and against each other. As expected, retroactive spies and retroactive criminals did not differ in face bias during encoding (*t*(74) = 0.17, *p* = .865), consistent with neither group knowing at the time of viewing which role they would later be assigned. Neither retroactive group differed significantly from control (retroactive spy vs. control: *t*(74) = 1.21, *p* = .232; retroactive criminal vs. control: *t*(74) = 1.06, *p* = .294; unadjusted planned comparisons), supporting their inclusion alongside neutral participants as the reference condition.

### 3.3. Improved responsiveness to encoding material hampered by personal goal state

The engagement task confirmed that across all conditions, participants became increasingly familiar with the encoding material, with participants able to detect more coins faster across blocks (detection: *χ*^2^(2) = 41.11, *p* < .001; speed: *F*(2, 17740.7) = 59.09, *p* < .001). Whilst there were no main effects of condition (detection: *χ*^2^(2) = 0.14, *p* = .932; speed: *F*(2, 94.9) = 0.94, *p* = .396), the presence of a goal state significantly altered whether participants got faster at responding (*F*(4, 17743.2) = 6.16, *p* < .001): Whilst proactive criminals and controls increased their response speed between blocks 1 and 3 (criminal: *z* = 6.68, *p* < .001; control: *z* = 10.86, *p* < .001), proactive spies were unable to do so (*z* = 1.61, *p* = .242). Potentially, the more rigid viewing behaviour in proactive spies described above (e.g., viewer fixations, shorter scanpath length) contributed to emerging differences to other conditions that viewed the scenes more thoroughly and could thus detect coins more readily.

It should be noted, though, that not all locations were equally accessible across the screen, with the semantic vicinity of the coin influencing both detection and speed heavily (detection: *χ*^2^(2) = 1151.42, *p* < .001; speed: *F*(2, 100.6) = 293.92, *p* < .001). The probability of detection was much lower and slower for coins near objects (61.8%, .56 s) than personal (83.6%, .49 s) or spatial features (85.6%, .52 s). Overall, participants detected more spatial than personal coins (*z* = 2.98, *p* = .008) but the pattern of detection according to categories did not differ across conditions (*χ*^2^(4) = 4.99, *p* = .288). Similarly, while condition interacted with response times across categories (*F*(4, 100.3) = 3.06, *p* = .020), this was not meaningful, since no condition differed significantly in response speed to each type of semantic vicinity, especially not the experimental conditions spy and criminal (*ps* > .210). Therefore, we can assume that participants across conditions engaged with the encoding material to a similar degree.

### 3.4. Narrative behaviour differed only in respect to goal categories

Given that goal state shaped gaze allocation in the naturalistic learning paradigm, we analysed whether these attention biases during encoding changed how participants accessed their resulting memory during free recall. Importantly, on this experimental day they had not received any reminder of their goal state.

Across all proactive conditions and the control group, we analysed how much of the participants’ speech was dedicated to the different scored semantic categories by calculating quasi-binomial general linear models on the percentages (Figure 4A). In line with our predictions, the only scored categories differing across groups were personal information (*χ*^2^(2) = 6.67, *p* = .036) and spatial information (*χ*^2^(2) = 7.80, *p* = .020). Planned contrasts confirmed that the spy and criminal group differed significantly in how much of their speech represented personal (*z* = 2.56, *p* = .010) and spatial content (*z* = −2.67, *p* = .008). We observed no differences in percentage of speech used to express insecurity (*χ*^2^(2) = .41, *p* = .815), unrelated information (*χ*^2^(2) = .19, *p* = .910), quoting verbal speech (*χ*^2^(2) = .89, *p* = .640), information on relating to the control category objects (*χ*^2^(2) = .58, *p* = .747) or other categories (*χ*^2^(2) = .09, *p* = .958).

**Figure 4:**
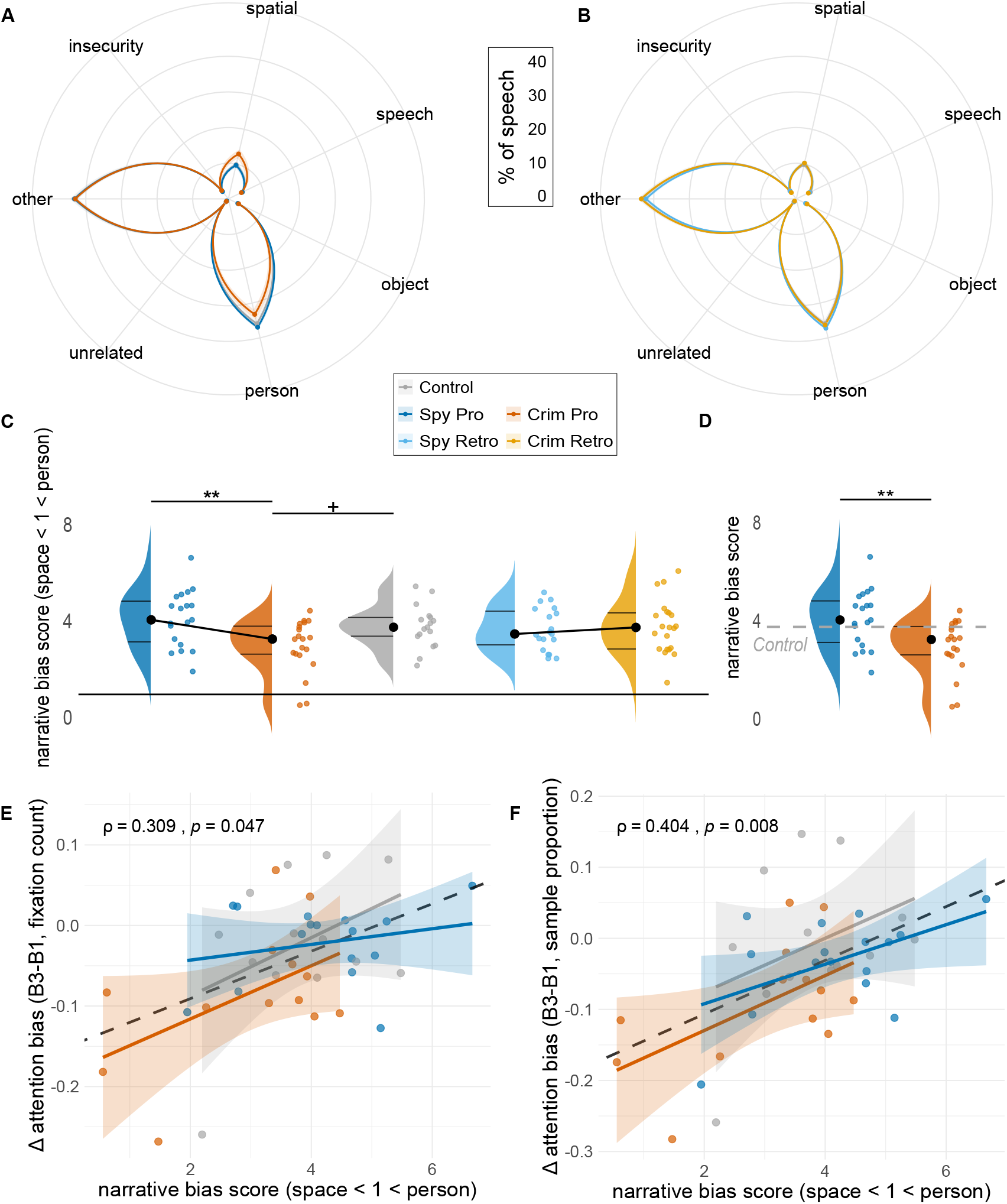
Goals modulate content of narrative recall. Radial plots for proactive (**A.**) and retroactive (**B.**) groups illustrate similar narrative structure across semantic categories apart from goal-relevant categories in proactive groups only: spies (blue) emphasising personal details over criminals (orange), and reverse for spatial details. Dots represent condition means, with shaded area as standard error of the mean. **C.** All narrative bias scores (n = 97) show person bias above the black reference line indicating no bias (1). Criminals show a reduction of bias in favour of spatial details against spies or control for proactive groups only. Black dots denote condition means, black lines quantiles. **D.** Proactive group difference from C. contrasted in reference to control mean (dotted grey line). Positive correlations between narrative bias scores, and change in attention bias drawn from eye-tracking fixation counts (**E.**, n = 42) and sample proportions (**F.**, n = 42). Change was calculated as attention bias in block 3 - block 1. Upregulating attention bias towards faces related to higher narrative bias scores, whilst lowering the attention bias in favour of places was associated with more place-oriented narrative bias scores. Black dashed line indicates overall Spearman correlation fit, colours of dots and shaded lines correspond to individual values and fitted lines for each condition.

Goal state thus only altered the speech profile in respect to the relevant conditions. Furthermore, a secondary negative binomial model revealed no differences in the total word count (*χ*^2^(2) = .25, *p* = .883), suggesting that goal-states did not change consistently how much participants chose to talk during free recall.

The respective analyses focused on retroactive conditions and the control group confirmed neither differences in the total word count across conditions (*χ*^2^(2) = .25, *p* = .884), nor any differences across the semantic categories contained within their verbal speech (insecurity: *χ*^2^(2) = .16, *p* = .923, unrelated information: *χ*^2^(2) = .27, *p* = .872, verbal speech: *χ*^2^(2) = .74, *p* = .691, objects: *χ*^2^(2) = 1.75, *p* = .416, spatial: *χ*^2^(2) = .15, *p* = .928, personal: *χ*^2^(2) = 1.76, *p* = .414, other: *χ*^2^(2) = 1.66, *p* = .435, Figure 4B).

### 3.5. Narrative content reflected proactive goal states

After confirming that participants’ narrative behaviour only differed with regard to the goal-related categories, we analysed whether the content of their episodic-like memory, accessed via free recall, was systematically influenced by their goal-state. A linear mixed model on proactive conditions and controls revealed that condition significantly affected the narrative bias score (*F*(2, 53) = 5.12, *p* = .009). Tukey-corrected post-hoc tests with robust standard errors confirmed that this difference was driven by a reduction of the narrative bias score indicating the recall of more spatial information in the criminal condition (*M* = 2.98, *S E* = .24) in contrast to the spy condition (*M* = 4.11, *S E* = .27; *t*(53) = 3.33, *p* = .0045, Figure 4C, D). The change in the bias score toward more spatial information in criminals in contrast to control (*M* = 3.81, *S E* = .21) reached trend levels (*t*(53) = 2.30, *p* = .064), whilst spy and control conditions did not differ significantly (*t*(53) = −0.92, *p* = .634). We ran an additional confirmatory analysis using a normalised bias score, calculated analogously to the attentional bias score above. The results corroborated the significant difference in the planned contrast between proactive spies and criminals (*t*(53) = 2.74, *p* = .0083), with a marginal overall effect of condition in the robust Wald test (*F*(2, 53) = 2.79, *p* = .0704), with criminals showing a reduced face bias (*M* = 0.437, *S E* = .058) in contrast to spies (*M* = 0.589, *S E* = .024). The pattern of results suggests that criminals adjusted the naturally occurring person bias towards place information in accordance with their space-focused goal state.

The same results emerged when considering only factually accurate narrative information: Proactive goal-state significantly altered the narrative bias score (*F*(2, 53) = 5.24, *p* = .008), with criminals showing the lowest bias score, thus towards spatial information (*M* = 2.98, *S E* = .24). Narrative bias scores in the criminal condition were significantly lower than in the spy (*M* = 4.11, *S E* = .27; *t*(53) = 3.31, *p* = .0047; Tukey corrected), and at trend level compared with the control condition (*M* = 3.80, *S E* = .22; *t*(53) = 2.33, *p* = .060; Tukey corrected). Confirmatory analyses using a normalised bias score showed a significant difference in the planned contrast between proactive spies and criminals (*t*(53) = 2.75, *p* = .0081, Wald test group model: *F*(2, 53) = 2.82, *p* = .068), with criminals showing a reduced face bias (*M* = 0.435, *S E* = .058) in contrast to spies (*M* = 0.587, *S E* = .024). There were much fewer data points when restricting the analysis to factually incorrect information, with no difference in narrative bias score emerging between conditions for neither the narrative bias score (*F*(2, 32) = 0.17, *p* = .846), nor the normalised variation (*F*(2, 32) = 0.03, *p* = .971).

Interestingly, the goal-state related shift in narrative bias score occurred despite all participants showing a strong narrative bias towards personal information, consistent with the biased viewing behaviour during encoding: One sample t-tests revealed a deviation of the narrative bias score towards personal as opposed to spatial information across and within all conditions (all: *t*(95) = 24.15, *p* < .001; pro spy: *t*(18) = 11.70, *p* < .001; pro criminal: *t*(19) = 8.21, *p* < .001; retro spy: *t*(18) = 13.3, *p* < .001; retro criminal: *t*(20) = 11.40, *p* < .001; control: *t*(16) = 13.20, *p* < .001, Figure 4C). This bias towards personal data was also observed when narrative content was divided by factual accuracy, showing a bias for accurately (*t*(95) = 23.96, *p* < .001) and wrongly recollected content (*t*(59) = 8.00, *p* < .001).

In contrast to the proactive conditions, a retroactively induced goal-state did not cause a condition-based shift in narrative bias (*F*(2, 54) = .11, *p* = .900, Figure 4C). There were also no condition differences when the narrative content was divided by accuracy (accurate: *F*(2, 54) = .08, *p* = .920, false: *F*(2, 29) = .70, *p* = .506).

Specifically, contrasting the factors of timing (pro vs retro) and job (spy vs criminal) indicated an overall effect of job (*F*(1, 75) = 4.22, *p* = .043), with spies showing higher face bias in the narrative bias score. There was no significant effect of timing (*F*(1, 75) = .67, *p* = .415), but a significant interaction (*F*(1, 75) = 6.65, *p* = .012), indicating that the effect of goal state depends on when the goal state was introduced. Beyond confirming that only proactive conditions showed an effect of goal-state (proactive: *t*(75) = 3.29, *p* = .0015; retroactive: *t*(75) = .34, *p* = .732), post-hoc analyses on the interaction also revealed that a proactive goal state shifted narrative bias in criminals towards spatial information (*t*(75) = −2.38, *p* = .0199) whereas spies displayed a strong bias towards personal information regardless of when the goal state was introduced (*t*(75) = 1.29, *p* = .202). This may indicate that spies continue to display a high natural social bias, whereas criminals, having a competing goal, actively shift this bias, particularly when the goal state is present already during memory encoding. This interpretation was further supported by the equivalent results when only considering factually accurate content (job: *F*(1, 75) = 4.56, *p* = .036, timing: *F*(1, 75) = .70, *p* = .404, interaction: *F*(1, 75) = 6.49, *p* = .013): Shifts in narrative bias scores were driven by an active prioritisation of spatial features in the proactive criminal condition specifically (*t*(75) = −2.37, *p* = .020), weakening the prevalent person bias that remains dominant in spies (*t*(75) = 1.25, *p* = .214).

### 3.6. Changes in attention viewing bias related to narrative content bias

We further analysed whether the extent to which goals modulated the attention bias during encoding was related to goal-specific regulation of the narrative bias. When averaged across all encoding blocks, none of the eye-tracking measures estimating attention bias correlated with the narrative bias for the proactive and the control conditions (count: *ρ* = .08, *p* = .630; duration: *ρ* = .03, *p* = .841; proportion: *ρ* = .20, *p* = .211; *n* = 42). However, the change in attention bias, measuring how much the goal state modulated the attention bias across encoding blocks, significantly correlated with narrative bias for all measurement types: An increased count of gaze allocations towards faces as opposed to spaces was linked to a higher narrative bias (*ρ* = .31, *p* = .047, Figure 4E). Furthermore, increased duration in face fixations (*ρ* = .35, *p* = .023), and an increased proportion of viewing time (*ρ* = .40, *p* = .008, Figure 4F) were associated with higher narrative bias scores. Therefore, the extent to which goal-state shaped attention bias over encoding was directly related to later narrative bias. There were no significant correlations between any measure of attention bias and narrative bias (*ps* > .510, *n* = 47), nor between the change in attention bias and narrative bias (*ps* > .598) when modelling all conditions without an active goal state during encoding (control, retroactive conditions). This indicates that active goal-states during an episodic-like task shape attentional allocation over time, and that this altered attentional allocation has lasting downstream effects on how participants remember these experiences.

## 4. Discussion

In this study, we aimed to trace how our individual goals shape the formation and content of realistic episodic-like memories by using naturalistic stimulus material and analysis methods. Specifically, we manipulated behavioural goal-states by introducing secret jobs, which motivated emphasis on personal (spies) or spatial (criminals) information. Indeed, our results show that these goal-states significantly modulated how complex scenes were perceived and later narrated. Viewing behaviour was naturally biased towards faces, but the bias decreased with increasing familiarity. Notably, this viewing trajectory was shaped by the goal-state, keeping spies more attuned to faces and allowing criminals to depreciate faces more strongly in favour of spatial information. Similarly, when freely narrating their episodic-like memories, participants showed a general bias toward person-related information. Nonetheless, a spatial goal-state allowed criminals to shift their narrative towards spatial content. Importantly, the adaptation of viewing bias across time was related to the strength of the narrative bias during memory recall. The strong bias towards faces or people across both gaze and narrative measures is consistent with the literature on the selective attention of humans to social stimuli (Langton et al., 2008; Skripkauskaite et al., 2023). Humans even show such preferential treatment for faces or people that these social stimuli constitute a distraction from the task, even if the visual input is optically matched for saliency with equivalent objects (Doherty et al., 2019). The naturally occurring social attention bias was likely additionally enhanced by the nature of the stimulus material, narrative films, used in our study. Both character-driven or plot-driven storytelling put the dominant attentional focus on people by following either one person, or the actions or social interactions of one or multiple actors. The results on viewing behaviour and task engagement support this interpretation: In general, proactive spies explored less within the film clips, showing fewer but longer-held fixations and decreased scanning paths, consistent with a motivated focus on more centrally placed goal-relevant social information. The reduced visual exploration in proactive spies can also account for the lack of increased engagement speed that was observed in the other conditions. The more fluid exploration shown by controls and especially proactive criminals driven by their spatially motivated goal state enabled them to engage with encoding material more speedily with increasing familiarity. Taken together, these results indicate a general viewing emphasis on people across groups, but especially strengthened in proactive spies whose goal-state highlighted persons. The strong shift of bias scores towards people was not only expected but a feature of the task design and validates the implemented bias measures, which detected the well-known social bias phenomenon.

Crucially though, whilst participants in our study showed a strong social attention bias, they were able to modify this bias according to their assigned goal state across encoding blocks. Despite initially similar viewing patterns, participants managed to shift the employment of their visual attention subtly towards their required goal dimension: Proactive spies maintained a strong person bias throughout the encoding task, while proactive criminals decreased their bias continuously in favour of spatial information. These findings replicated the literature for goal-directed attention in static stimuli (Itier et al., 2007; Vogt et al., 2010) but extended the observation to moving narrative stimuli. Even within such complex changing environments, participants adapted their gaze behaviour with increasing expertise. The attentional draw from faces decreased in the control group over repeated exposure to the stimulus material, suggesting that participants were able to explore the scene more freely when they became more familiar with the material. These results matched previous observations that social bias remained robust against task demands (Palanica and Itier, 2012), but was modulated by familiarity, with infrequent facial stimuli drawing bigger oculomotor responses (Pereira et al., 2022).

The goal state also shaped how participants remembered the naturalistic stimulus material. Participants modulated their narrative bias according to their respective goal: Whilst person-related information was dominant across all conditions, narrative bias was shifted to favour spatial information for criminals. Importantly, condition effects were exclusively driven by a modified emphasis on personal or spatial details, since the remaining narrative structure, such as length or signs of insecurity, did not differ across participants. This suggests that the attention manipulation had no general impact on the extent or detail of narrative recall but rather specifically modulated the content, biasing it towards goal-directed information. Crucially, the narrative shift by goal state also held true when only considering factually accurate information, speaking to the relevance of considering what was actually presented when analysing naturalistic memory. Our results therefore corroborated the classical finding of goals providing a biased encoding framework for the memory, which remains apparent during narrative recall (Pichert and Anderson, 1977), while extending this finding to naturalistic film stimuli. Importantly, the narrative bias must have occurred due to an underlying difference in the mnemonic representation itself, rather than as a result of biased recall due to task instructions, since participants did not receive a reminder of their goal state on the retrieval day. A potential mechanistic interpretation is that proactive goals provide a top-down schematic framework during memory encoding, restructuring or selectively tagging content for consolidation, which is then more accessible during later recall. Further research would be required to disentangle whether the narrative bias was driven by a differential storage of the memory itself or whether goal-related elements were associated with increased accessibility and thus verbalised more readily during speech production (Dell, 1986; Levelt, 1989). Nonetheless, the idea of a biased encoding framework was supported by the relationship of attentional and narrative biases: The extent to which the top-down goal state modified the naturally occurring social attention bias across encoding correlated positively with the narrative bias during retrieval. To rephrase, if a participant maintained a person bias across encoding, they showed a stronger person bias during narrative recall, and if they depreciated faces during encoding in favour of spatial information, they showed a stronger shift towards spatial information during recall. Therefore, in our results, not only were gaze fixation locations predictive of their memories (Celli et al., 2025), but so was the goal-directed alignment of these gaze locations. The observation that the magnitude of this alignment correlated with later mnemonic bias strongly supports the idea of memory being encoded within a biased framework that is increasingly aligned with the goal-relevant schemata.

The absence of retroactive goal effects highlights the relevance of a biased encoding framework: We did not observe differential narrative biases if the goal state was introduced only after the memory encoding, such that encoding did not occur under a goal-specific framework. Potentially, a schematic framing is necessary to tag goal-relevant content for preferential processing during consolidation. Consistent with this hypothesis, we did not observe a relationship between attentional and narrative bias in the retroactive condition. This mirrored findings in the literature emphasising the necessity of goal states present during encoding to predict memory (Pazzaglia et al., 2014), but contrasted with literature findings of effective retroactive biases (Patil et al., 2017; Braun et al., 2018). Potentially, the retroactive bias was not strong enough to affect the interconnected complexity of naturalistic memories. Mechanistically, prioritising stability in the representation of episodic memories might be preferable to prevent these complex memories from drifting and adapting too much too fast. Alternatively, a narrative bias score might simply not be sensitive enough to detect more subtle shaping from a retroactive bias. There might be other narrative markers indicative of the underlying memory representation, such as syntactic realisation (Pokhoday et al., 2019), that were not analysed in this study. Further research would be required to evaluate whether retroactive biases can exert any influence on naturalistic stimuli, for example with different behavioural measures or analysing whether post-encoding consolidation showed preferential processing of goal-relevant dimensions.

We applied a novel methodological approach using naturalistic measures to analyse the rich episodic-like memories. We utilised advances in computer vision to automatically annotate the film clips by semantic categories, thereby allowing the statistical analysis of the content of the complex visual input and interaction with it. Prior to the advent of such semantic segmentation models, it would have been highly costly to match and then analyse visual gaze behaviour within moving films. Using computer vision-based segmentation opens many possibilities of analysing naturalistic viewing behaviour as well as resulting mnemonic traces. Within the current study, the linguistic analysis of the free narrative content demonstrated a promising avenue to assess the underlying structure of complex, episodic-like memories. We were able to detect nuanced differences in retrieval based on the associated goal states that were introduced just the day before.

In conclusion, we found that proactive goals shaped how participants interacted with naturalistic film material, influencing the development of their visual attention during encoding and the narrative content of their recall. The extent to which attentional goals modulated their visual gaze related to goal-specific biases in their memory, suggesting that our attentional bias frames the resulting memory in line with our goals. The analyses of visual gaze behaviour with semantic segmentation models and the data-driven content decoding of the narrative recall preserved the rich, nuanced content of the data and represent promising avenues for future research.

## 5. Additional Information

### 5.1. Author contributions

HH: Methodology, Software, Investigation, Data Curation, Formal Analysis, Writing - Original Draft and Review & Editing, Visualisation; MM: Data Curation, Investigation, Formal Analysis; MS: Conceptualisation, Resources, Writing - Review & Editing, Funding acquisition; NMR: Conceptualisation, Methodology, Software, Formal analysis, Investigation, Data Curation, Writing - Original Draft and Review & Editing, Visualisation, Supervision, Project administration, Funding acquisition.

### 5.2. Data and Code Availability

The code is available on GitHub, including the statistical analyses. Both the eye-tracking and behavioural data can be found on Zenodo.

### 5.3. Acknowledgements

We thank Hanna Scharte, Jule Bemetz, Laura Hertel, Julius Debertshäuser, Jonathan Schmidt, Anisoara Diaconu, Anja Kobler, and Enna Mocnik for their help with data collection. This work was supported by funding from the German Research Foundation (HH: Project-ID 499552394 – CRC 1597 “SmallData”; NR: RO6828/1-1, RO6828/1–2; MS: SCHO1820/2–1) and the BMBF.

## Appendix A. Supplementary Methodology

### Appendix A.1. Eye-tracking analysis pipeline

**Figure A.5:**
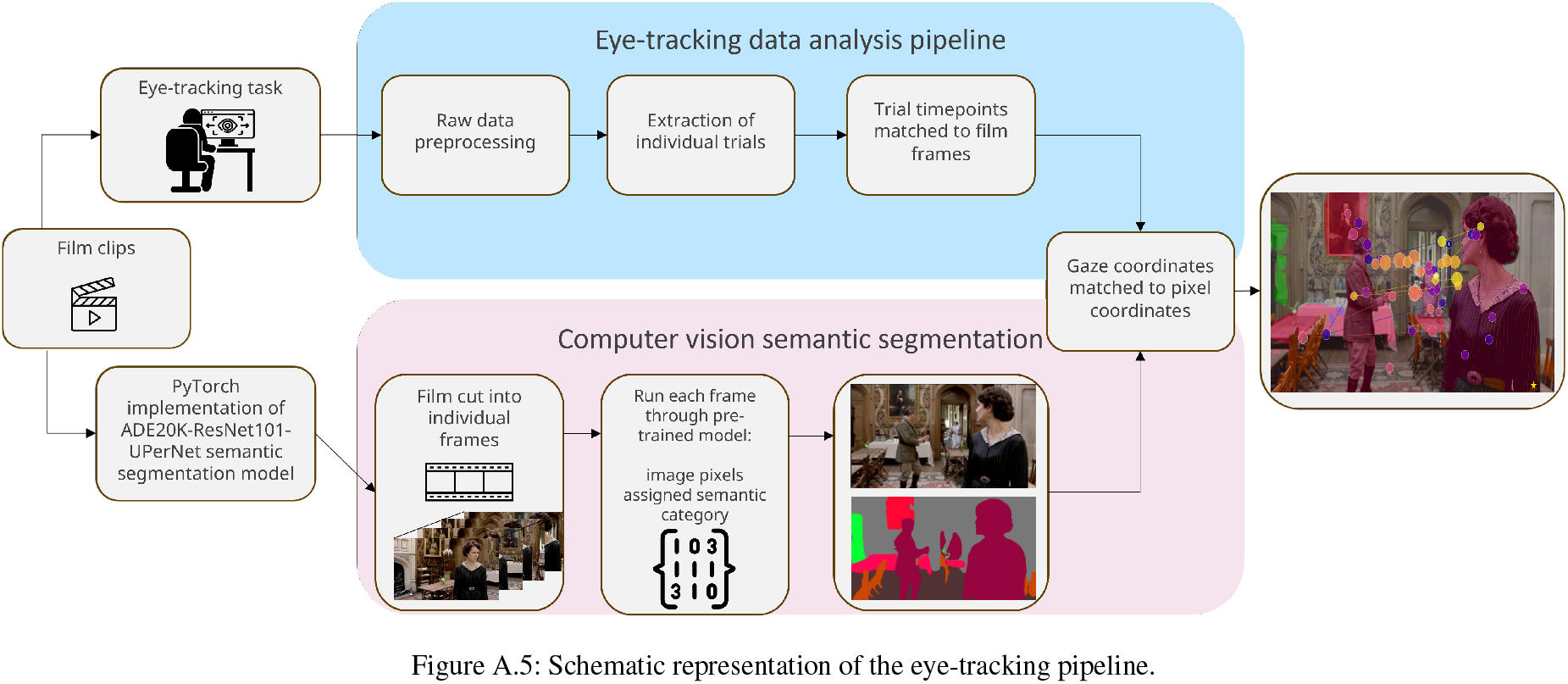
Schematic representation of the eye-tracking pipeline.

### Appendix A.2. Free-recall narrative category descriptions

**Table A.1:**
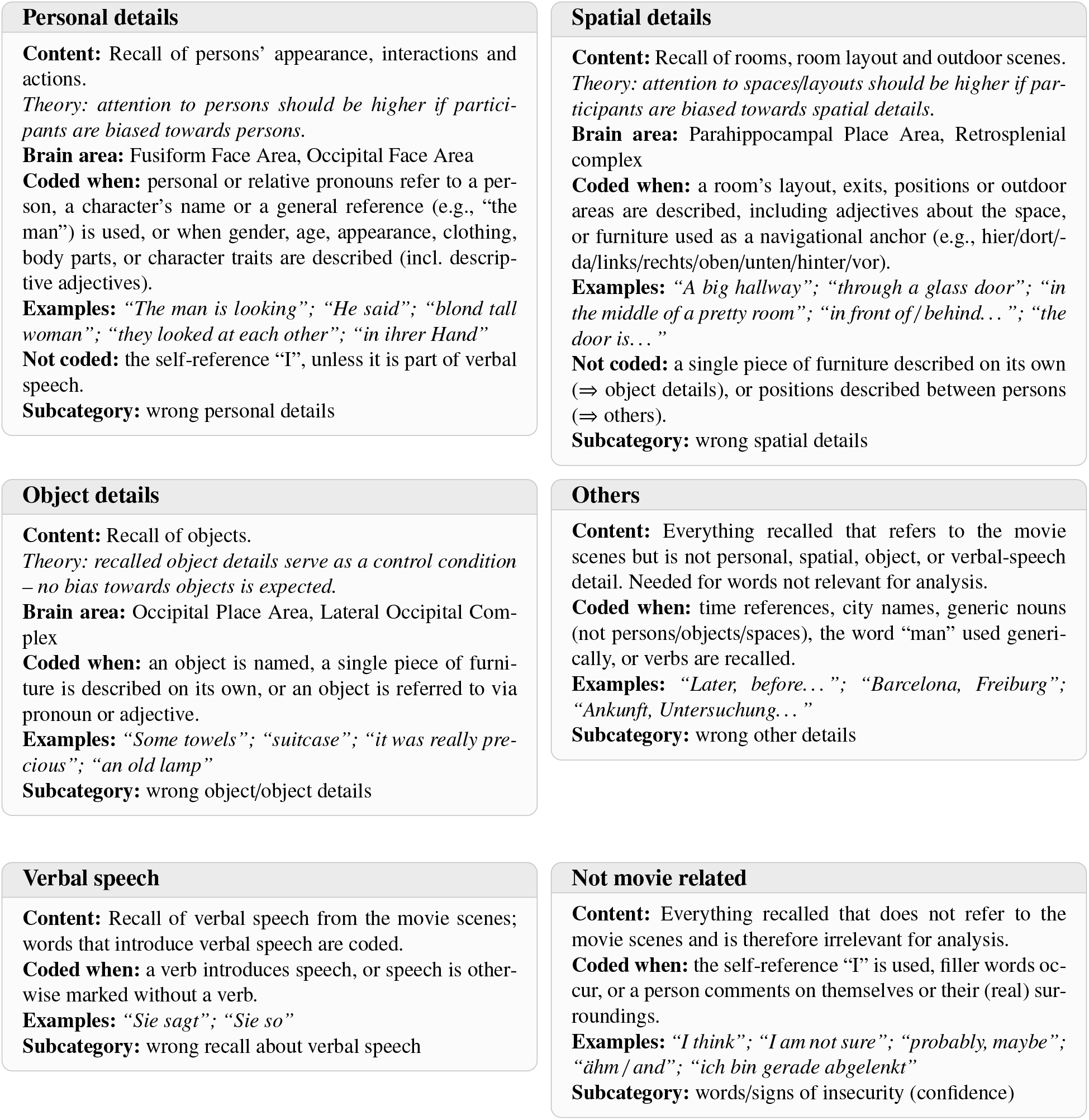
Coding scheme for movie scene recall categories (Rädiker and Kuckartz, 2019)

| Personal details | Spatial details |
|---|---|
| **Content:** Recall of persons’ appearance, interactions and actions.<br>*Theory: attention to persons should be higher if partici-pants are biased towards persons.*<br>**Brain area:** Fusiform Face Area, Occipital Face Area<br>**Coded when:** personal or relative pronouns refer to a per-son, a character’s name or a general reference (e.g., “the man”) is used, or when gender, age, appearance, clothing, body parts, or character traits are described (incl. descrip-tive adjectives).<br>**Examples:** *“The man is looking”; “He said”; “blond tall woman”; “they looked at each other”; “in ihrer Hand”*<br>**Not coded:** the self-reference “I”, unless it is part of verbal speech.<br>**Subcategory:** wrong personal details | **Content:** Recall of rooms, room layout and outdoor scenes.<br>*Theory: attention to spaces/layouts should be higher if par-ticipants are biased towards spatial details.*<br>**Brain area:** Parahippocampal Place Area, Retrosplenial complex<br>**Coded when:** a room’s layout, exits, positions or outdoor areas are described, including adjectives about the space, or furniture used as a navigational anchor (e.g., hier/dort/-da/links/rechts/oben/unten/hinter/vor).<br>**Examples:** *“A big hallway”; “through a glass door”; “in the middle of a pretty room”; “in front of / behind…”; “the door is…”*<br>**Not coded:** a single piece of furniture described on its own (⇒ object details), or positions described between persons (⇒ others).<br>**Subcategory:** wrong spatial details |
| **Object details** | **Others** |
| **Content:** Recall of objects.<br>*Theory: recalled object details serve as a control condition – no bias towards objects is expected.*<br>**Brain area:** Occipital Place Area, Lateral Occipital Com-plex<br>**Coded when:** an object is named, a single piece of furni-ture is described on its own, or an object is referred to via pronoun or adjective.<br>**Examples:** *“Some towels”; “suitcase”; “it was really pre-cious”; “an old lamp”*<br>**Subcategory:** wrong object/object details | **Content:** Everything recalled that refers to the movie scenes but is not personal, spatial, object, or verbal-speech detail. Needed for words not relevant for analysis.<br>**Coded when:** time references, city names, generic nouns (not persons/objects/spaces), the word “man” used generi-cally, or verbs are recalled.<br>**Examples:** *“Later, before…”; “Barcelona, Freiburg”; “Ankunft, Untersuchung…”*<br>**Subcategory:** wrong other details |
| **Verbal speech** | **Not movie related** |
| **Content:** Recall of verbal speech from the movie scenes; words that introduce verbal speech are coded.<br>**Coded when:** a verb introduces speech, or speech is other-wise marked without a verb.<br>**Examples:** *“Sie sagt”; “Sie so”*<br>**Subcategory:** wrong recall about verbal speech | **Content:** Everything recalled that does not refer to the movie scenes and is therefore irrelevant for analysis.<br>**Coded when:** the self-reference “I” is used, filler words oc-cur, or a person comments on themselves or their (real) sur-roundings.<br>**Examples:** *“I think”; “I am not sure”; “probably, maybe”; “ähm / and”; “ich bin gerade abgelenkt”*<br>**Subcategory:** words/signs of insecurity (confidence) |

## References

Aly, M., Turk-Browne, N.B., 2016. Attention promotes episodic encoding by stabilizing hippocampal representations. Proceedings of the National Academy of Sciences 113, E420–E429.

Bindemann, M., Burton, A.M., Hooge, I.T., Jenkins, R., De Haan, E.H., 2005. Faces retain attention. Psychonomic bulletin & review 12, 1048–1053.

Braun, E.K., Wimmer, G.E., Shohamy, D., 2018. Retroactive and graded prioritization of memory by reward. Nature communications 9, 4886.

Celli, M., Cona, G., Volpi, T., Tronelli, V., Zangrossi, A., Corbetta, M., 2025. Predicting visual object memory through natural eye movement topography. Scientific Reports.

Cerf, M., Frady, E.P., Koch, C., 2009. Faces and text attract gaze independent of the task: Experimental data and computer model. Journal of vision 9, 10–10.

Cowan, N., Bao, C., Bishop-Chrzanowski, B.M., Costa, A.N., Greene, N.R., Guitard, D., Li, C., Musich, M.L., Übal, Z.E., 2024. The relation between attention and memory. Annual review of psychology 75, 183–214.

Damiano, C., Walther, D.B., 2019. Distinct roles of eye movements during memory encoding and retrieval. Cognition 184, 119–129.

Dell, G.S., 1986. A spreading-activation theory of retrieval in sentence production. Psychological review 93 3, 283–321. URL: https://api.semanticscholar.org/CorpusID:21975212.

Doherty, B.R., Fraser, A., Nobre, A.C., Scerif, G., 2019. The functional consequences of social attention on memory precision and on memory-guided orienting in development. Developmental Cognitive Neuroscience 36, 100625.

Duchowski, A.T., Duchowski, A.T., 2017. Eye tracking methodology: Theory and practice. SpringerLink.

Dunsmoor, J.E., Murty, V.P., Davachi, L., Phelps, E.A., 2015. Emotional learning selectively and retroactively strengthens memories for related events. Nature 520, 345–348.

van Ede, F., Nobre, A.C., 2023. Turning attention inside out: How working memory serves behavior. Annual review of psychology 74, 137–165.

End, A., Gamer, M., 2019. Task instructions can accelerate the early preference for social features in naturalistic scenes. Royal Society open science 6, 180596.

Fehlmann, B., Coynel, D., Schicktanz, N., Milnik, A., Gschwind, L., Hofmann, P., Papassotiropoulos, A., de Quervain, D.J., 2020. Visual exploration at higher fixation frequency increases subsequent memory recall. Cerebral cortex communications 1, tgaa032.

Itier, R.J., Villate, C., Ryan, J.D., 2007. Eyes always attract attention but gaze orienting is task-dependent: Evidence from eye movement monitoring. Neuropsychologia 45, 1019–1028.

Jonikaitis, D., Moore, T., 2019. The interdependence of attention, working memory and gaze control: behavior and neural circuitry. Current opinion in psychology 29, 126–134.

Knowlton, B.J., Castel, A.D., 2022. Memory and reward-based learning: A value-directed remembering perspective. Annual review of psychology 73, 25–52.

Kowler, E., 2011. Eye movements: The past 25 years. Vision research 51, 1457–1483.

Langton, S.R., Law, A.S., Burton, A.M., Schweinberger, S.R., 2008. Attention capture by faces. Cognition 107, 330–342.

Levelt, W.J.M., 1989. Speaking: From Intention to Articulation. The MIT Press. URL: 10.7551/mitpress/6393.001.0001, doi:10.7551/mitpress/6393.001.0001.

Madore, K.P., Wagner, A.D., 2022. Readiness to remember: predicting variability in episodic memory. Trends in cognitive sciences 26, 707–723.

Palanica, A., Itier, R.J., 2012. Attention capture by direct gaze is robust to context and task demands. Journal of Nonverbal Behavior 36, 123–134.

Patil, A., Murty, V.P., Dunsmoor, J.E., Phelps, E.A., Davachi, L., 2017. Reward retroactively enhances memory consolidation for related items. Learning & memory 24, 65–69.

Pazzaglia, A.M., Staub, A., Rotello, C.M., 2014. Encoding time and the mirror effect in recognition memory: Evidence from eyetracking. Journal of Memory and Language 75, 77–92. URL: https://www.sciencedirect.com/science/article/pii/S0749596X14000588, 10.1016/j.jml.2014.05.009.

Pereira, E.J., Birmingham, E., Ristic, J., 2022. Infrequent faces bias social attention differently in manual and oculomotor measures. Attention, Perception, & Psychophysics 84, 829–842.

Pichert, J.W., Anderson, R.C., 1977. Taking different perspectives on a story. Journal of educational psychology 69, 309.

Pokhoday, M., Shtyrov, Y., Myachykov, A., 2019. Effects of visual priming and event orientation on word order choice in russian sentence production. Frontiers in Psychology 10, 1661.

Rayner, K., 1998. Eye movements in reading and information processing: 20 years of research. Psychological bulletin 124, 372.

Ryan, J.D., Shen, K., 2020. The eyes are a window into memory. Current opinion in behavioral sciences 32, 1–6.

Rädiker, S., Kuckartz, U., 2019. Analyse qualitativer Daten mit MAXQDA. Springer.

Sabo, M., Wascher, E., Schneider, D., 2024. Neural basis of attentional selection in goal-directed memory retrieval. Scientific reports 14.

Schmidig, F.J., Yamin, D., Sharon, O., Nadu, Y., Nir, J., Ranganath, C., Nir, Y., 2025. Anticipatory eye gaze as a marker of memory. Communications psychology 3.

Skripkauskaite, S., Mihai, I., Koldewyn, K., 2023. Attentional bias towards social interactions during viewing of naturalistic scenes. Quarterly Journal of Experimental Psychology 76, 2303–2311.

Song, H., Finn, E.S., Rosenberg, M.D., 2021. Neural signatures of attentional engagement during narratives and its consequences for event memory. Proceedings of the National Academy of Sciences 118, e2021905118. URL: https://www.pnas.org/doi/abs/10.1073/pnas.2021905118, doi:10.1073/pnas.2021905118, arXiv:https://www.pnas.org/doi/pdf/10.1073/pnas.2021905118.

Sterpenich, V., van Schie, M.K., Catsiyannis, M., Ramyead, A., Perrig, S., Yang, H.D., Van De Ville, D., Schwartz, S., 2021. Reward biases spontaneous neural reactivation during sleep. Nature communications 12, 4162.

Sulewski, P., Amme, C., Hebart, M.N., König, P., Kietzmann, T.C., 2026. Fixation duration on natural scenes is explained by memory encoding not processing demand. Nature neuroscience 29, 1488–1497.

Trajkovska, K., Kljun, M., Pucihar, K.Č., 2024. Gaze2aoi: Open source deep-learning based system for automatic area of interest annotation with eye tracking data. arXiv preprint arXiv:2411.13346.

Vogt, J., De Houwer, J., Moors, A., Van Damme, S., Crombez, G., 2010. The automatic orienting of attention to goal-relevant stimuli. Acta psychologica 134, 61–69.

Zhou, B., Zhao, H., Puig, X., Fidler, S., Barriuso, A., Torralba, A., 2017. Scene parsing through ade20k dataset, in: Proceedings of the IEEE Conference on Computer Vision and Pattern Recognition.

Zhou, B., Zhao, H., Puig, X., Xiao, T., Fidler, S., Barriuso, A., Torralba, A., 2018. Semantic understanding of scenes through the ade20k dataset. International Journal on Computer Vision.

